# Quantitative high-resolution imaging of live microbial cells at high hydrostatic pressure

**DOI:** 10.1101/847228

**Authors:** A. C. Bourges, A. Lazarev, N. Declerck, K. L. Rogers, C. A. Royer

## Abstract

The majority of the Earth’s microbial biomass exists in the Deep Biosphere, in the deep ocean and within the Earth’s crust. While other physical parameters in these environments, such as temperature or pH, can differ substantially, they are all under high-pressures. Beyond emerging genomic information, little is known about the molecular mechanisms underlying the ability of these organisms to survive and grow at pressures that can reach over 1000-fold pressure on the Earth’s surface. The mechanisms of pressure adaptation are also important to in food safety, with the increasing use of high-pressure food processing. Advanced imaging represents an important tool for exploring microbial adaptation and response to environmental changes. Here we describe implementation of a high-pressure sample chamber with a 2-photon scanning microscope system allowing for the first time, quantitative high-resolution two-photon imaging at 100 MPa of living microbes from all three kingdoms of life. We adapted this setup for Fluorescence Lifetime Imaging Microscopy with Phasor analysis (FLIM/Phasor) and investigated metabolic responses to pressure of live cells from mesophilic yeast and bacterial strains, as well as the piezophilic archaeon, *Archaeoglobus fulgidus*. We also monitored by fluorescence intensity fluctuation-based methods (scanning Number and Brightness (sN&B) and Raster scanning Imaging Correlation Spectroscopy (RICS)) the effect of pressure on the chromosome-associated protein HU and on the ParB partition protein in *E. coli*, revealing partially reversible dissociation of ParB foci and concomitant nucleoid condensation.

**SIGNIFICANCE:** The majority of the Earth’s microbial biomass exists in high-pressure environments where pressures can reach over 100 MPa. The molecular mechanisms that allow microbes to flourish under such extreme conditions remain to be discovered. The high pressure, high resolution imaging system presented here revealed pressure dependent changes in metabolism and protein interactions in live microbial cells, demonstrating great promise for understanding deep life.

## INTRODUCTION

Life on Earth exists in a wide range of environmental conditions including extremes of pressure, temperature, pH or salt concentrations (1). Indeed, over 90% of the Earth’s microbial biomass is thought to exist in high-pressure environments (2). Pressures encountered by living organisms on Earth can reach >100 MPa in the deepest ocean trench (the Challenger Deep in Mariana trench) or within the Earth’s crust. Several dozen microbial isolates from the deep biosphere have been cultured, and a few of their genomes have been sequenced (3), but these represent only a small fraction of the diversity of species anticipated for this ecosystem. To date, the molecular mechanisms that allow these organisms to survive and proliferate at high pressure are not understood. Progress in this area would have implications for understanding the emergence of life on Earth, mechanisms that control carbon cycling and the search for life elsewhere. High pressure food processing is today a 14 billion-dollar industry and is expected to quadruple in the next decade (4). Acquired resistance to pressure treatment by foodborne pathogens represents a serious economic issue (5), the mitigation of which will require insight into the molecular mechanisms of microbial adaptation and response to pressure.

Direct observation at high pressure of organisms from the deep biosphere and their atmospheric pressure adapted counterparts using advanced microscopy would yield important information about these mechanisms. However, the poor pressure resistance of sample holder materials has required HP microscopy cells with thick windows, severely limiting optical resolution (6). Here we present the adaptation of a fused silica capillary system, first used for *in vitro* Fluorescence Correlation Spectroscopy (7, 8) to high-pressure, high resolution fluorescence imaging. We have implemented improvements in sample loading, adherence and washing in order to apply such a system to the study of live microbial cells using quantitative high-resolution imaging techniques directly under pressure.

We report here the first high resolution quantitative observations by two-photon microscopy of molecular processes taking place in live microbial cells under high hydrostatic pressure. We examined the pressure-response of microbes from all three kingdoms of life, two adapted to atmospheric pressure, the Gram-positive bacterium, *Escherichia coli*, and the yeast, *Saccharomyces cerevisiae*, as well as the anaerobic piezophilic thermophilic archaeon, *Archaeoglobus fulgidus*, a sulfur-metabolizing organism found in hydrothermal vents. The natural auto-fluorescence from metabolic enzyme cofactors (NAD(P)H and FAD) (9) represents an alternative live cell imaging contrast to fluorescent proteins. We coupled two-photon excitation of NAD(P)H (740 nm) with scanning Fluorescence Lifetime Imaging Microscopy (FLIM) and Phasor analysis (10). FLIM/Phasor analysis of NAD(P)H fluorescence provides a metabolic footprint of live bacterial cells (11). In *E. coli,* we compared total NAD(P)H to the metabolic state before, during and after pressurization. These studies revealed a strong and complex response of *E. coli* metabolism to pressure.

In cases for which fluorescent proteins can be genetically introduced into the sample strains, quantitative imaging modalities that rely on the fluorescence intensity fluctuations such as scanning Number and Brightness (sN&B) (12) and Raster Imaging Correlation Scanning (RICS) (13) analyses can be applied. These approaches yield protein stoichiometry, absolute concentration, and diffusion properties (12, 13). Here we applied fluorescence fluctuation-based methods, sN&B and RICS, to monitor the effect of pressure on the stoichiometry, localization and dynamics of two fluorescent protein fusions implicated in nucleoid structure and plasmid partitioning and co-expressed in *E. coli* cells. Our results revealed pressure-induced dissociation of partition complexes and pressure-induced foci formation of the nucleoid-associated protein. These pressure effects were reversible for only a subset of cells.

## MATERIALS AND METHODS

### Cell preparation for imaging

*Escherichia coli* K12 reference strain MG1655 and its *mrr*^−^ derivative devoid of the Mrr endonuclease were used for FLIM experiments. For sN&B measurements, the *E. coli* strain DLT3053 was used for constitutive co-expression of the HU-mCherry fusion encoded by a chromosomal insertion and ParB-mVenus fusion encoded by the plasmid pJYB234 (14). The protocol was similar to that described previously (15). Overnight cultures were diluted in LB medium and grown at 37°C until the OD_600_ reached 0.6. The cells were centrifuged and resuspended in a few μL to a final OD_600_ ~25 in minimal M9 medium supplemented with 0.4% glucose. *Saccharomyces cerevisiae* BY4741 (MATa; his3Δ 1; leu2Δ 0; met15Δ 0; ura3Δ) was grown on full SC medium agar plate and then a single colony was used to set an overnight culture in SC medium supplemented by 2% glucose and diluted the next morning. Similar to bacteria, a high density of cells was required for the injection in the capillary. *Archaeoglobus fulgidus* was grown in a liquid heterotrophic (lactate) sulfate-reduction medium using anoxic techniques ((16) and references therein). A cell pellet of an overnight culture was resuspended, under anoxic conditions, to obtain a highly-concentrated cell suspension prior to injection. A few μL of the highly concentrated cell preparation were injected into the coated capillary. The cells were left for a few minutes (~10 min) to attach to the surface. Those that were not attached were rinsed with the fresh appropriate medium (M9 minimal medium for bacteria, SC medium for yeast and media as described previously (16) for the archaea used to purge the entire system in order to prevent cells death.

### Capillary preparation and coating

The middle of the 15-inch capillary was burned with a lighter for 2 to 3 seconds to remove the outer polyimide coating. The capillary was passed through the two glands (Fig. 1A) and glued to the drilled pressure plug using epoxy glue. A solution of 100x chitosan for immobilization (17) was freshly prepared with approximately 0.015 g of chitosan powder diluted in 900 μL of 2 M (= 10%) glacial acetic acid in an Eppendorf tube. Then, 60μl of this 100x solution were diluted in 900 μl of distilled water. The chitosan solution was passed into the capillary using the peristatic pump until the solution was apparent in the output tubing and the connector. Incubation was allowed for 20-30 min at room temperature. The chitosan solution was rinsed by flushing the system with the minimal medium required by the cells during the experiment.

**FIGURE 1.**
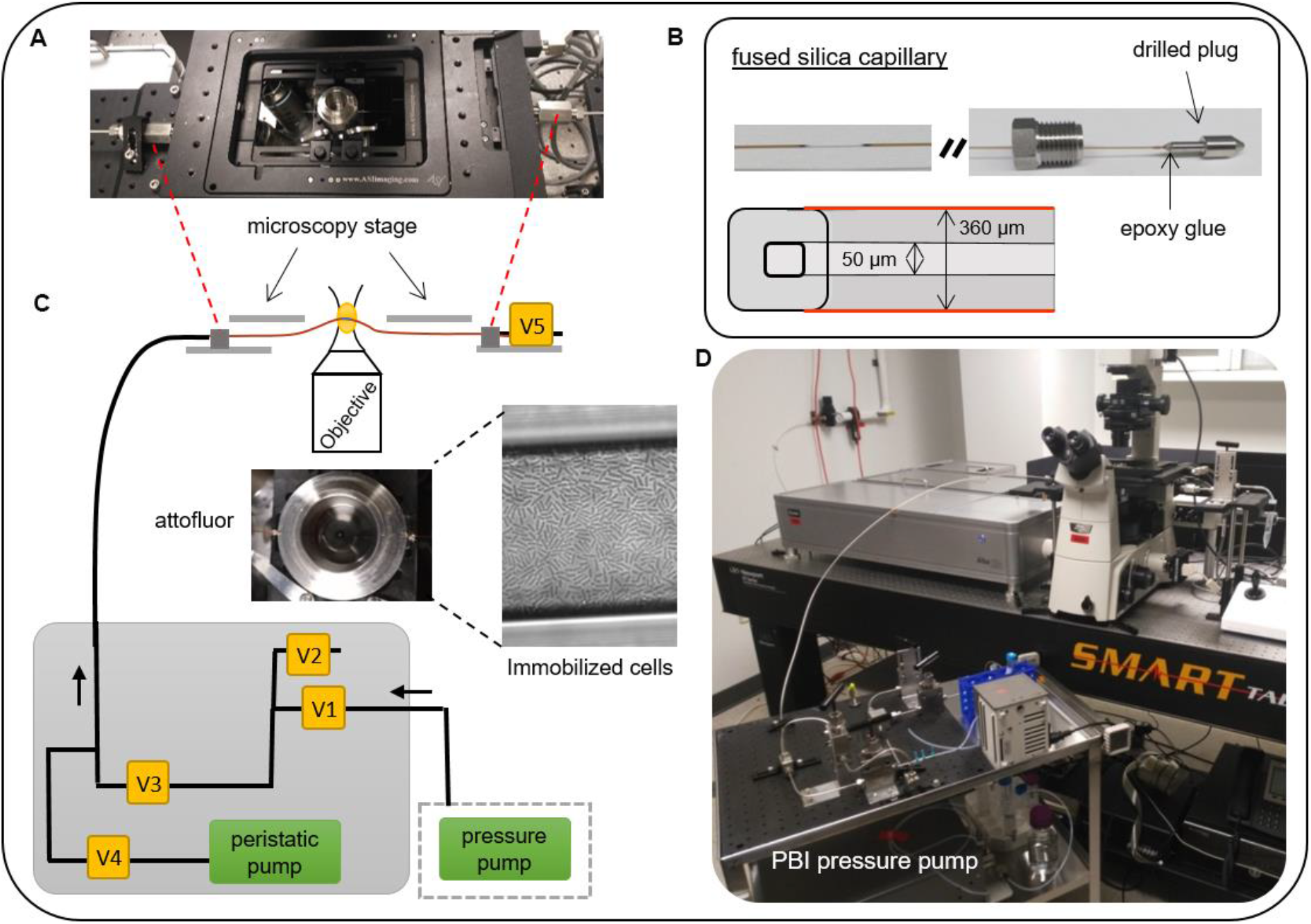
Microscope setup used to perform high resolution quantitative imaging of live cells under pressure. (A) Microscopy stage showing the capillary immobilized in a holder (attofluor) between a coverslip and the objective with glycerol as a coupling media. (B) cross-section and photo of the fused silica circular capillary with an external polyimide coating inserted and glued in drilled plugs. (C) Schematic of the HP connections. Two valves (V) make it possible to switch with either the peristatic pump or the pressure pump to load or apply pressure, respectively (V3 and V4). (D) Photograph of the cart (with pressure pump, lines and valves as designated in the schematic, and the connection to the microscope with the mounted capillary.

### Sample loading

The capillary was moved into the modified attofluor holder, which was placed on the microscope stage so that the burned portion of the capillary was located at the objective. The attofluor is slitted at 180 degrees such that the capillary is held in place. The 15-inch length of the capillary allows for easy adjustment of the capillary position on the microscope stage. The capillary was seated on a rubber gasket at the bottom of the attofluor to avoid breakage, and a second rubber gasket was placed on top of the capillary. Then, the capillary was immobilized in the stainless-steel holder with 2 coverslips (N2, VWR) on the top to prevent bending upon contact with the objective, and a stainless-steel gasket was placed within the ring of the attofluor. The weight of this piece maintains the capillary z-position during focusing. Glycerol was used as coupling medium. By closing V3 and opening V4, the system was connected to the peristatic pump. A drop (less than 10 μl) of concentrated solution of cells (OD_600_ ~25) was deposited on the plug located at the end of the capillary near V5 and the peristaltic pump was run backwards until cells passing through the capillary were visible on the camera or at the objective. The pump was then turned off and the capillary tightly connected to the pressure output valve, V5. The cells were allowed to attach to the surface for 10 minutes, and then with V4 closed and V3 and V5 open, unattached cells were rinsed away with minimal medium supplemented with 0.4% glucose for *E. coli* cells and the growth medium noted above for the yeast and archaea at a low flow rate from the pressure pump (0.05 ml/min). This method allowed a flat field of view with a single layer of immobilized cells that remained attach to the surface with increasing pressure (Fig. S1). When there were no more unattached cells (Fig. S1), the pressure pump was switched off and V5 closed, leaving the system ready for high pressure experiments.

All pressure connections (tubing, lines, glands) were 1/8 in in diameter (High Pressure Equipment, Inc., Erie PA) to minimize the footprint of the high-pressure apparatus. Moreover, we used an RF-1700 constant high-pressure pump from (Pressure BioSciences, Inc., South Easton, MA) with a maximum pressure set to 15,000 psi (~100 MPa). In order to match the fused silica capillary refraction index, oil was replaced by glycerol as the coupling medium. All tubing and pumps were purged with minimal media to minimize background fluorescence. Pressure was increased using a low flow rate (0.05 μl/min) such that reaching 100 MPa required 2-3 minutes. Pressure was released by slowly opening the output valve.

### Fluorescent Lifetime Imaging Microscopy with Phasor Analysis (FLIM/Phasor)

The advantage of this phasor approach for FLIM analysis lies in the easy interpretation of the raw data without any fitting of lifetime decay curves (10). Each pixel of the FLIM image was transformed into a pixel on the phasor plot with *g* and *s* coordinates calculated from the fluorescence intensity decay using the following transformation with *i* and *j* corresponding to a pixel of the image and ω the frequency determined by the laser repetition rate (80 MHz):

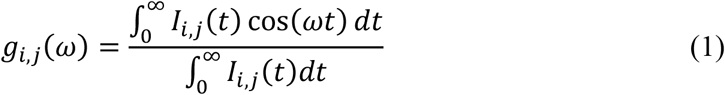

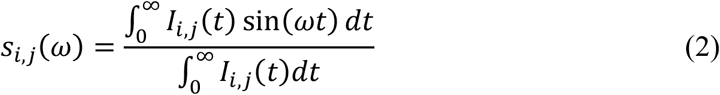

Pixels corresponding to a single species with a single exponential fluorescence intensity decay are located on the universal semicircle limiting the phasor plot and the position on the semi-circle depends on the lifetime. Short lifetimes are located on the right with high *g* values and small *s* values, while long lifetimes are near the origin of the semicircle. Multi-exponential decays are located inside the semicircle, at a position defined by the lifetime values and their relative fractional intensities. For a mixture of two single exponential decays, the phasor will be located on a line between the two-single decay positions on the universal plot, the position being weighted by the fractional contribution of each single exponential component to the decay. Indeed, the coordinates *g* and *s* are described following by equations 3 and 4 with *h*_*k*_ the intensity and *τ*_*k*_ the lifetime of component, *k*:

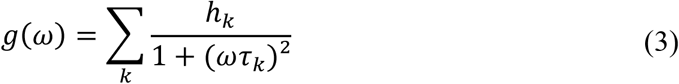

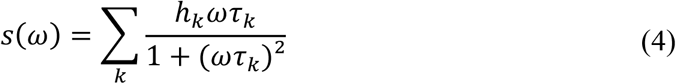

Thus, the global phasor at each pixel of an image is the sum of the independent phasors of each fluorescent decay. To highlight the spatial localization of lifetime components, we selected a cluster of pixels within the phasor plot and visualized the localization of these pixels on the fluorescent image using the VistaVision software (ISS, Champaign, IL, USA). FLIM experiments were performed at 740 nm with a two-photon excitation and a frequency of 80 MHz. The number of frames, the pixel dwell-time and the laser power were adjusted in order to collect 100 counts in the brightness pixel of the field of view (FOV). We have generally acquired a single frame with 1 ms of pixel dwell-time for an FOV of 13 ×13 μm and 256 ×256 pixels.

### Number and Brightness (N&B)

Number and Brightness (N&B) is an image-based implementation of fluctuation spectroscopy (12). Here we imaged fluorescent protein fusions expressed in bacteria. A series of raster scans (25 frames in our case) was acquired using a pixel dwell-time (40 μs) faster than the diffusion time. This provides fluorescence intensity values over time for each pixel from which fluorescence fluctuations (variance) and average intensity (*F*) can be calculated and used to deconvolve the fluorescence intensity into the brightness (*B*) and the number of molecules (*N*) in the excitation volume (*F=N x B*). The shot noise corrected brightness (*e*) values and the number (*n*) of diffusing molecules were calculated at each pixel:

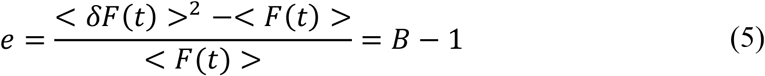

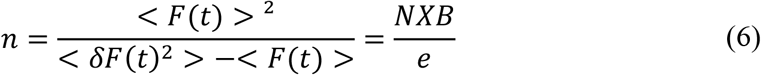

The brightness is directly proportional to the number of fluorophore-containing molecules diffusing together, so that the brightness of a monomeric fluorescent protein can be used to calculate the stoichiometry of an oligomeric complex. However, the auto-fluorescence of the background contribution tends to decrease the values of brightness (*esample*) and fluorescence intensity (*Fsample*) measured, which is why we have removed its contribution using N&B measurements on a background strain that does not express FP-tagged fluorescence molecules (brightness *ebg* and fluorescence *Fbg*) following the equation 7:

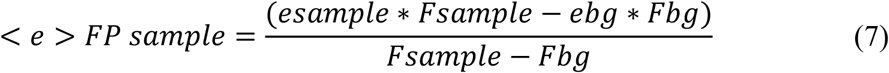

These analyses were carried out using the Patrack software (18) (Patrick Dosset, CBS, Montpellier, France). The average values were calculated for all pixels within the central region of all cells in the image as previously described for N&B studies on bacteria (19).

Scanning N&B was performed as previously described by Bourges et al. (15) except that an FOV was scanned 25 times instead of 50 times to control the time of exposure to pressure. The size of an FOV was 13X13 μm. All experiments on fluorescent protein fusions were performed with a 2-photon excitation at 930 nm. For the simultaneous excitation of mVenus and mCherry, a 580LP mirror was used to separate the emission light on two channels with a 530/43 nm (mVenus) or 650/50 nm (mCherry) filter (Chroma Technologies). Merged images of the average fluorescence intensity images were obtained using the software package, Fiji (available online at http://fiji.sc/). The background corrected brightness values presented were calculated using the background strain *E. coli* MG1655 (15).

### Raster Imaging Correlation Spectroscopy (RICS)

From the same raster scans obtained in N&B, the diffusion coefficient of the fluorescent molecules can be extracted by fitting the pixel pair spatio-temporal correlation function (20),

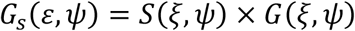

where *ξ* and *ψ* are the spatial increments of the scanning laser in *x* and *y*, respectively. The *S*(*ξ, ψ*) and *G*(*ξ, ψ*) functions correspond to the correlation due to scanning and that due to diffusion, respectively

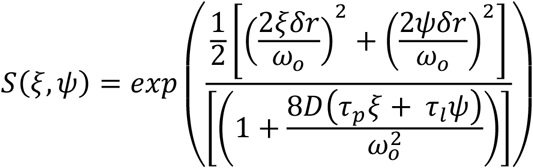

and

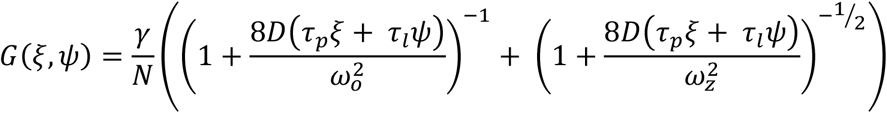

where *τ*_*p*_ and *τ*_*l*_ are the pixel dwell time and line time, respectively, and *δr* is the distance between pixels in a line.

The pixel dwell-time used was 40 μs, which means that one frame in sN&B took 2-3 s. Thus, fluorescence fluctuations were observed even for particles diffusing very slowly on the millisecond time scale. The vertical (line to line) spatio-temporal correlation function is on the millisecond timescale, and is thus, the most useful. It reflects the fact that the probability of seeing a molecule in the next line is higher if the molecule diffuses slowly. Conversely, if the molecule diffuses quickly, such as free diffusing GFP, the fluorescence intensity correlation decreases rapidly from line to line and is represented by a rapid drop in the vertical autocorrelation curve. Raster scans were converted to RICS curves and analyzed using SimFCS (E. Gratton, LFD, Irvine, CA, USA).

## RESULTS AND DISCUSSION

### Live cell imaging compatible high-pressure sample chamber

The major modifications presented here with respect to prior high-pressure single molecule *in vitro* setups involved implementing a sample loading system that was compatible with immobilizing and washing microbial cells and scaling down the size of the high-pressure components as described in the Methods section. We employed square capillary tubing as previously described (21), which while presenting lower pressure limits (~150 MPa) has significantly better optical properties and allows for efficient immobilization of microbial cells. The thickness of the capillary walls (150 μm) matches the working distance of high N.A. objectives. First, rather than sealing one end of the capillary with a blow torch after sample loading (7, 21) (a process that would be highly detrimental to live cells), a drilled plug system sealed around the capillary with epoxy glue was used for both ends (Methods and Fig 1B). Thus, both ends of the capillary were glued into high-pressure plugs that were drilled with a 400 mm bore. Because both ends were connected to the pressure system via the drilled plugs, a peristaltic pump could be used to easily load the microbial cells into the capillary (Fig. 1B, Fig. S1). After loading the cells and washing out unadhered cells with medium, the capillary was connected to the pressure lines by switching between two valves, V3 and V4 (Fig. 1B). Pressure experiments were performed by closing a valve located at the output of the capillary (V5 on Fig. 1B). The 2-photon scanning microscope (Alba, ISS, Champaign Illinois) was the same as previously describe in Bourges et al. (15), with the exception of the objective that has been replaced by an 60X 1.4 NA oil immersion objective (Nikon APO, VC) (Fig. 1).

### Microbial metabolic responses to pressure

Measurements of Fluorescence Lifetime with Phasor analysis of the enzyme co-factors, NADH or NADPH provides a fingerprint of microbial metabolic states (11). The bound co-factors exhibit a significantly longer fluorescence lifetime than the free forms, although NADH and NAD(P)H cannot be distinguished from each other. The difference in fluorescence lifetimes for bound and free NAD(P)H results in distinct positioning on the phasor plot. As described in the Methods section, the *g* and *s* coordinates in the phasor plot represent, respectively, the cosine and sine Fourier transforms of the intensity decay function. Single exponential decays will have *s* and *g* phasor values such that they lie on the universal half circle (Fig. S2), with long lifetimes towards the left and short lifetimes toward the right. For samples with emitting species exhibiting different lifetimes, the phasor position will be defined by the fractional contribution of each of the species. In an image, the phasor position of each pixel within a field of view (FOV) is uniquely defined by the fluorescence decay at that pixel and the acquisition frequency (here 80 MHz). Pixels with a mixture of bound and free NAD(P)H will be situated below the universal circle at a position between the bound and free forms that depends upon their ratio. Thus, metabolic changes affect the position of the phasor position of pixels for images of NAD(P)H auto-fluorescence (9). Bacterial response to antibiotic stress is associated with higher ratios of free/bound NAD(P)H (11). For example, in *B. subtilis*, the switch from glycolysis to gluconeogenesis led to a shift toward a higher free/bound NAD(P)H ratio (Fig. S2), reflecting a change from catabolic to anabolic activity.

The pressure response of the NAD(P)H lifetime for three different cell types, *Escherichia coli, Archaeoglobus fulgidus (type strain VC-16)* and *Saccharomyces cerevisiae* (bacteria, archaea and yeast) was tested in the high-pressure capillary microscope system (Fig. 2). *Archaeoglobus fulgidus* is ubiquitous in subsurface, high-pressure environments and has been reported to grow up to 60 MPa (16). It is also an anaerobic thermophile (~83°C maximum growth temperature) (22), while the yeast and bacterium are both mesophilic organisms that grow aerobically at atmospheric pressure. We maintained anaerobic conditions during the experiment with *A. fulgidus* by purging the entire system with oxygen-depleted minimal medium prior to loading the cells.

**FIGURE 2.**
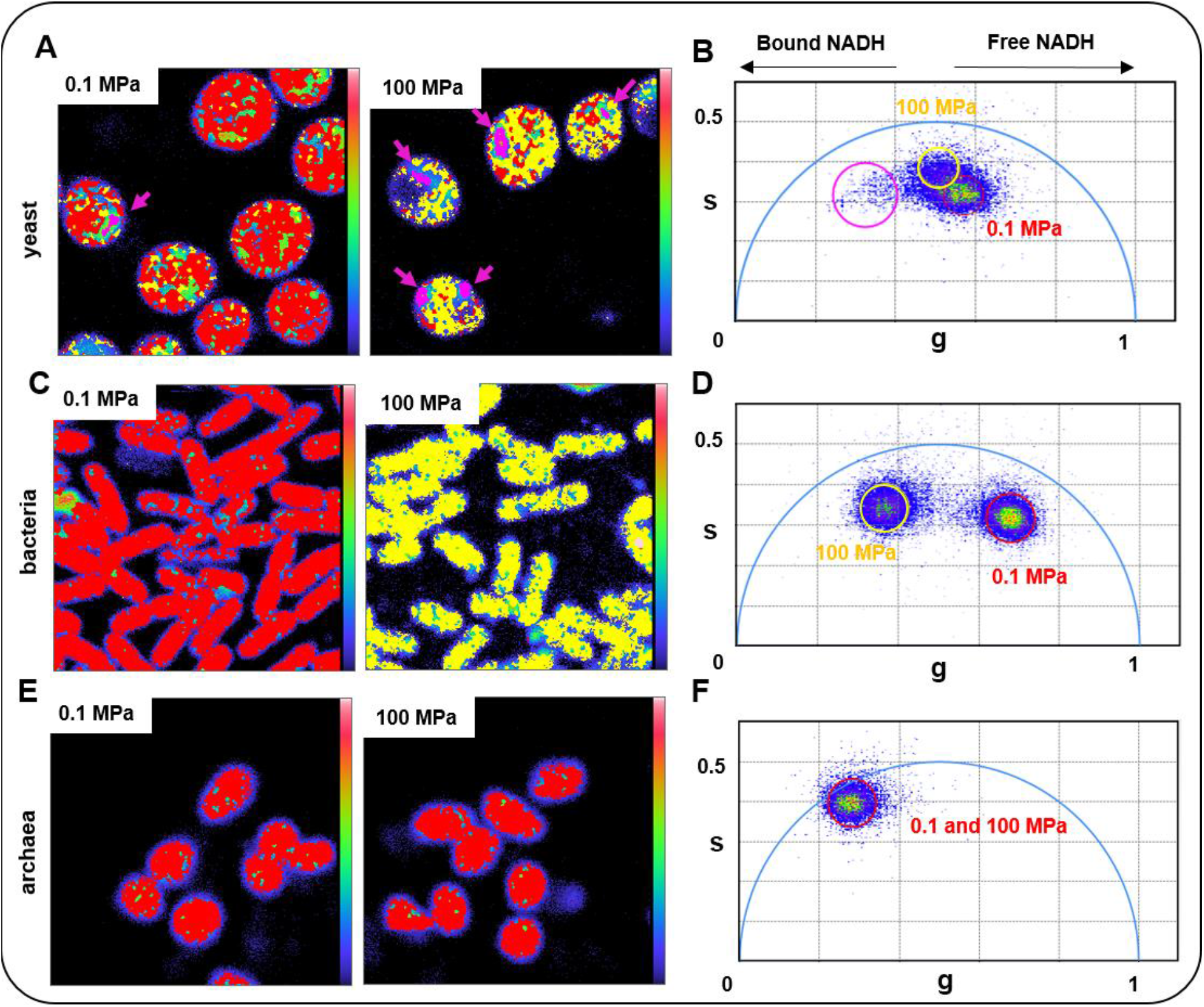
Effect of pressure on FLIM/Phasor of NADH for yeast (*Saccharomyces cerevisiae*), bacteria (*E. coli* MG1655) and archaea (*Archaeoglobus fulgidus*). (A, C and E) are fluorescent images with pixels colored according to their position on the phasor plots (B, D and F), respectively. Pixels are colored according to their phasor positions. Red 0.1 MPa and yellow and pink 100 MPa. Top row; *S. cerevisiae*, pink arrows correspond to foci with high bound/free NAD(P)H ratios. Middle row: *E. coli*, Bottom row: *A. fulgidus*. For *A. fulgidus*, at 100 MPa most pixels remained within the red circle at atmospheric pressure. Images correspond to a field of view of 13 ×13 μm for yeast and bacteria and 10 ×10 μm for archaea and 256 ×256 pixels. Excitation was 740 nm, and Phasor frequency was 80 MHz. At least 3 fields of view were acquired at each pressure point and the experiment was repeated 3 times with the same observations. Color scales reflect overall intensity, but intensity is obscured by the pixel phasor colors.

At atmospheric pressure, most pixels for the yeast cells were positioned at *g* values > 0.5, indicating a significant fraction of free NAD(P)H (Figs. 2A, B, red). At 100 MPa, most pixels in the yeast cells shifted slightly toward bound NAD(P)H (yellow), while some pixels (Fig. 2A, B, magenta) shifted strongly toward bound NAD(P)H species and were localized in small foci, indicating association of metabolic enzymes into discrete foci upon high-pressure stress. In contrast to the strong pressure response of yeast, we found no pressure response of the auto-fluorescence of the piezophile, *A. fulgidus* (Figs. 2e, f). Even at atmospheric pressure, the phasor values for its NAD(P)H auto-fluorescence were shifted strongly toward bound NAD(P)H, and this did not change at 100 MPa, even though the pressure used was higher than its maximum survival pressure (80 MPa) (16). It should be noted that *A. fulgidus* analyses were conducted at room temperature, well below the optimal growth range for this thermophilic species. Thus, the absence of a response to elevated pressures might be explained by the cells undergoing temperature stress well before pressurization. We are currently developing protocols to implement temperature control in the high-pressure imaging system.

The response of *E. coli* to pressure was stronger than for the other two microbes (Figs. 2C, D). At atmospheric pressure compared to the yeast sample, the average phasor position of the pixels was shifted towards free NAD(P)H (red), indicative of lower metabolic activity, perhaps due to the switch to minimal medium (Fig. 2C-left, D, red circle and pixels). Note that the exact position and distribution of the atmospheric phasor values varied somewhat between experiments. However, in all experiments at 100 MPa, a significant fraction of pixels of NAD(P)H fluorescence in *E. coli* exhibited a shift of their phasor positions to lower *g* values (*e.g.*, Fig. 2C-right, D, yellow circle and pixels), indicating a pressure-induced increase in the fraction of bound NAD(P)H (9).

The MG1655 reference strain of *E. coli* used in these experiments harbored a pressure-activated restriction endonuclease, Mrr, which leads to a pressure-induced SOS response (15, 23, 24). To ascertain whether response of *E. coli* metabolism to pressure was linked to this Mrr-induced SOS response, we carried out FLIM/Phasor analysis on the auto-fluorescence of an *E. coli* strain in which the *mrr* gene had been deleted *(mrr−).* Comparison of the average intensity images with the Phasor maps of an *E. coli* strain at atmospheric pressure shows that regardless of the overall NAD(P)H content (total intensity), most pixels exhibited similar fractions of bound/free co-factor (Fig. 3 A-C). These were somewhat shifted to lower *g*-values, perhaps reflecting differences in the growth rate for this sample. At 100 MPa pressure (Fig. 3D-F), in contrast to the *mrr*+ strain, a bi-stable pressure response of NAD(P)H lifetimes was observed. Comparison of the average intensity (correlated to the concentration of NAD(P)H) with the phasor patterns revealed that bacterial cells exhibiting a higher fraction of bound NAD(P)H (Fig. 3E, green pixels, left shifted) also exhibited a much lower overall auto-fluorescence intensity (Fig. 3D, green arrow). We conclude that these bacteria were compromised by high-pressure and that much of their free NAD(P)H had diffused out of the cells. Since the enzyme-bound fraction was less likely to diffuse out of these cells due to its larger molecular weight, the fraction of bound NAD(P)H increased in these cells. In contrast, cells that maintained high auto-fluorescent intensity were those with NAD(P)H phasor values shifted to the right towards free NAD(P)H (Fig. 3D-F, yellow pixels and arrow). This indicates decreased metabolic activity in response to pressurization. Upon return to atmospheric pressure, cells exhibiting low auto-fluorescence intensity mostly retained the left-shifted phasor pixels (Fig. 3G-I, green arrows and pixels), indicating that they had been irreversibly compromised by pressure. Some of the cells that retained their high auto-fluorescence intensity, also retained the right-shifted pixels corresponding to lower metabolic activity, (Fig 3G-I, yellow pixels and arrows). In contrast, in some cells with high auto-fluorescence intensity, the pixels recovered their original phasor positions (Fig. 3G-I, red pixels and arrows), indicating complete reversibility of the pressure effects on metabolic state for these cells. These FLIM/Phasor results highlight the stochastic nature and complexity of the response of bacterial metabolism to pressure. They establish that phenomena other than the SOS response participate in pressure-induced loss of cell viability.

**FIGURE 3.**
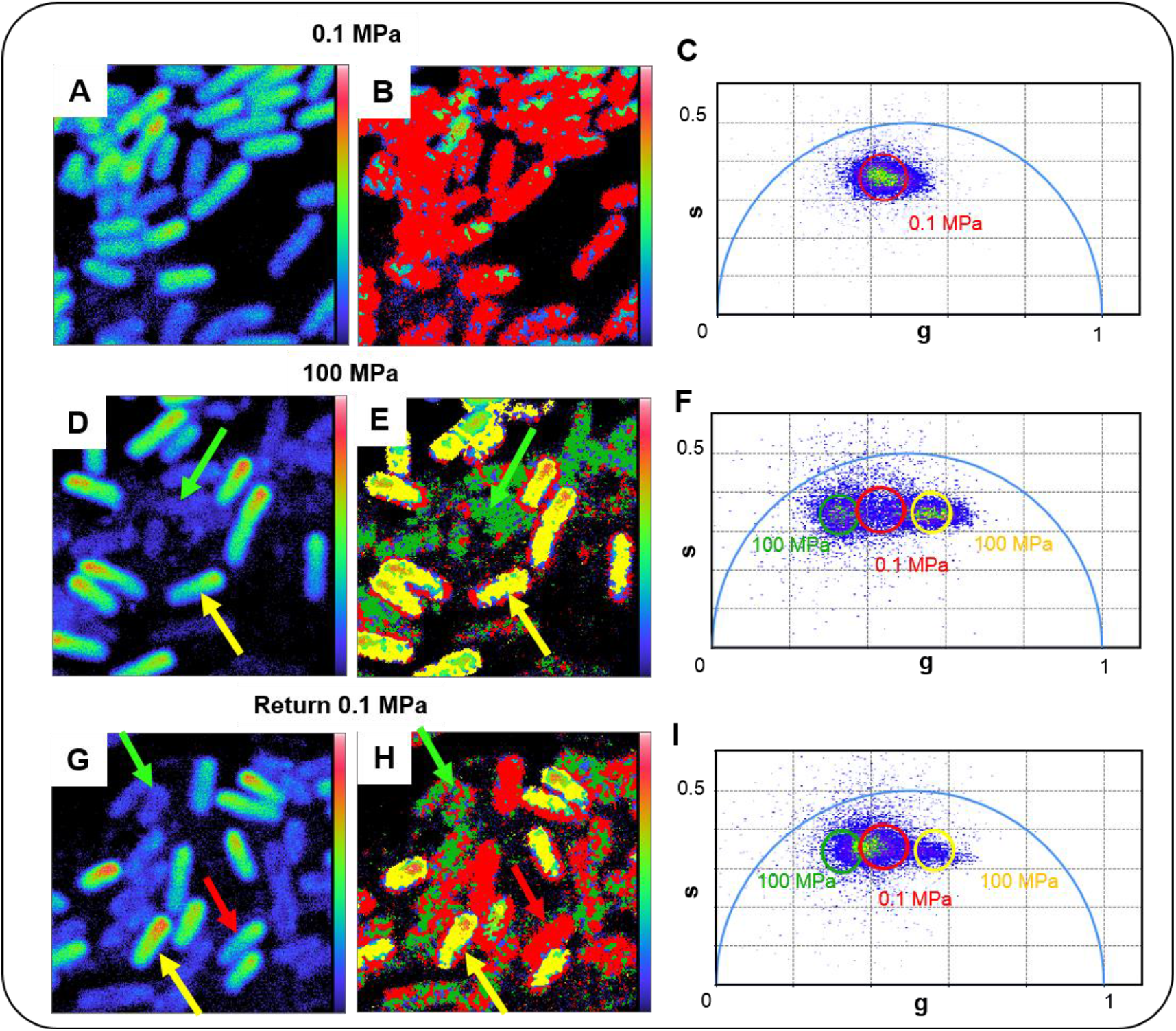
Comparison of the pressure response of total auto-fluorescence intensity with FLIM/Phasor of NAD(P)H for E. coli MG1655 (*mrr−*). (A-C) Atmospheric pressure. Fluorescence intensity, phasor map and phasor plots respectively. The red circle in C corresponds to the phasor positions of the red pixels in B. (D-F) 100 MPa Fluorescence intensity, phasor map and phasor plots respectively. The red, green and yellow circles in F correspond to the phasor positions of the red, green and yellow pixels in E respectively. The green arrows indicate a cell with low auto-fluorescence intensity and left-shifted phasor positions, while the yellow arrows indicate a cell with high auto-fluorescence intensity and right-shifted phasor values. Red pixels in the phasor plot represent positions that are unchanged with respect to atmospheric pressure. (G-I) Return to 0.1 MPa (atmospheric pressure). Fluorescence intensity, phasor map and phasor plots respectively. The red, green and yellow circles in I correspond to the phasor positions of the red, green and yellow pixels in H respectively. Green arrows indicate a cell with low auto-fluorescence intensity and left-shifted phasor positions, while yellow arrows indicate a cell with high auto-fluorescence intensity and right-shifted phasor values. Red arrows represent a cell with high intensity and phasor positions that are equivalent to those at atmospheric pressure. Images correspond to a field of view of 13 ×13 μm and 256 ×256 pixels. Excitation was 740 nm, and Phasor frequency was 80 MHz. At least 3 fields of view were acquired at each pressure point.

### >Pressure effects on the bacterial partition machinery

Large scale transposon mutagenesis on a moderate piezophile, *Photobacterium profundum,* revealed that the largest fraction of loci conferring pressure sensitivity to this organism were implicated in chromosome structure and function (25). This underscores the importance of proteins involved in chromosome function in pressure adaptation. Hence, we were interested in ascertaining the effects of pressure on bacterial proteins involved in such processes. As an example of such a system, we characterized the effects of pressure on the ParB partition complex protein of *E. coli* using our high pressure capillary microscopy system coupled with scanning Number and Brightness (sN&B) (12) and Raster Scanning Image Correlation Spectroscopy (RICS) (13). In *E. coli* the type I partition systems are involved in segregation of the F and P1 low copy number plasmids (26). The ParB protein recognizes *parS* sites on the plasmid and recruits the ParA ATPase. ParB is known to form dimers organized into foci (1 per plasmid origin of replication) containing ~200 ParB monomers (27, 28). These foci are thought to correspond to liquid-liquid phase separated droplets (14).

To monitor pressure effects on ParB in live *E. coli* cells we employed a strain in which ParB, fused to monomeric mVenus fluorescent protein, was expressed from its natural locus on the F plasmid (14). This strain expressed as well, a chromosomal mCherry fusion of the nucleoid associated protein, HU, to allow visualization of the entire nucleoid. At atmospheric pressure, discrete ParB-mVenus foci were apparent (Fig. 4A, green), while the HU-mCherry protein (Fig. 4A, red) was distributed homogeneously on the nucleoid. Application of 100 MPa pressure led to disruption of the ParB clusters (Figure 4B, green). In contrast, pressure caused the HU protein to form foci in most cells (Fig. 4B, red). Since this *E. coli* strain also expressed Mrr, a pressure-dependent restriction endonuclease (15, 23, 24), the formation of HU foci probably corresponded to DNA condensation after double-strand breaks and induction of the SOS response by Mrr (24). After pressure release, some bacterial cells appeared to recover, as evidenced by the reappearance of ParB-mVenus foci (Fig. 4C, green) and the dispersion of the HU-mCherry signal (Fig. 4C, red), while others did not.

**FIGURE 4.**
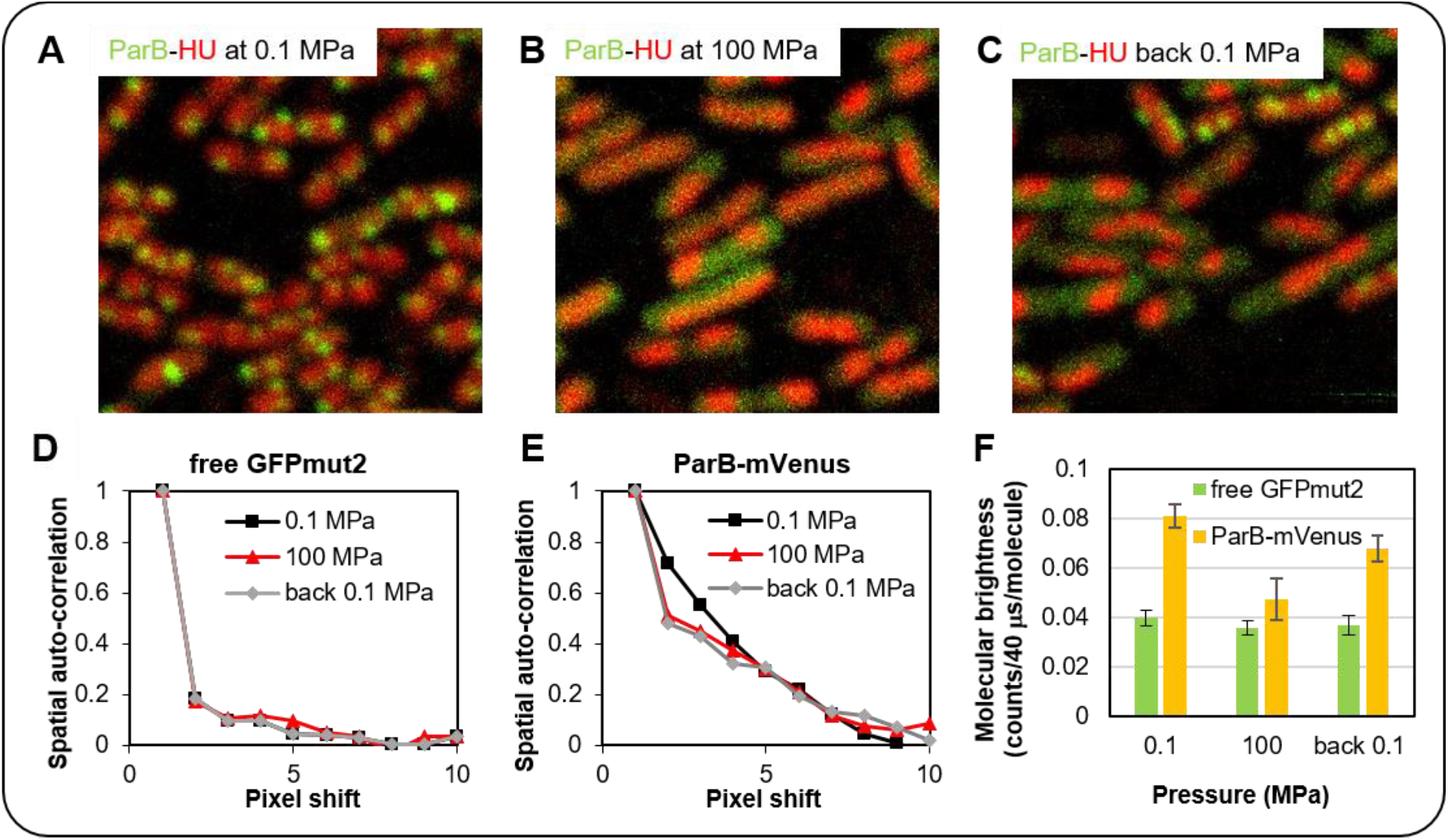
Effect of pressure on ParB Partition protein clusters and on the chromosome in *E. coli*. (A-C) Images correspond to the merging of the fluorescence intensity images of ParB-mVenus (in green) and HU-mCherry (in red) at (A) atmospheric pressure (0.1 MPa), (B) under pressure (100 MPa) and (C) back to atmospheric pressure (0.1 MPa). Images are 13 × 13 μm and 256 × 256 pixels. (D) and (E) Comparison of the vertical auto-correlation profile from RICS analyzes at 0 MPa (black squares), 100 MPa (red triangles) and back to 0.1 MPa (grey diamonds) of (D) freely diffusing GFPmut2 in E. coli cytoplasm and (E) mVenus fusion with ParB proteins in E coli. (F) Molecular brightness from the sN&B analyzes of free diffusing GFPmut2 (green) and ParB-mVenus (yellow) at atmospheric pressure (0.1 MPa), high pressure (100 MPa) and after pressure is released (back 0 MPa). GFPmut2 is expressed in E. coli MG1655 chromosome from the inducible *P_BAD_* promotor with 0.4% arabinose and HU-mCherry/ParB-mVenus protein fusions are constitutively expressed in *E. coli* DLT3053 with the plasmid pJYB234. The experiment was repeated 2 times with 8 FOV. Each RICS vertical auto-correlation curve is the average vertical auto-correlation profile of 8 FOV. The brightness values correspond to the average brightness values of the pixels inside bacteria of 16 FOV per pressure point with an average of 20 to 25 bacteria per FOV. Brightness values from all FOV from two days of experiments were averaged and corrected for background as described in Methods. Error bars represent the standard deviation of the mean for all 16 FOV.

RICS analysis of an *E. coli* MG1655 strain expressing free GFPmut2, a fast-maturing GFP variant (29), from a plasmid revealed that pressure did not affect the diffusion of free monomeric GFPmut2 (Fig. 4D). This is an interesting observation, as it indicates that the viscosity of the cytosol does not change significantly at 100 MPa pressure. In contrast, the average ParB-mVenus diffusion was faster under pressure and after release (Fig. 4E), consistent with dissociation of the foci at high pressure and limited recovery upon return to 0.1 MPa. In addition, the molecular brightness of mVenus fused to ParB calculated by sN&B decreased two-fold under pressure indicating that pressure led to the dissociation of ParB-mVenus dimers to monomers (Fig. 4F). Note that the molecular brightness of free GFPmut2 is not significantly affected by pressure *in vivo*. Thus, pressure appeared to dissociate both the ParB dimers and the foci, themselves, to form monomers that diffuse freely in the cytoplasm. As noted, it has been suggested that the ParB foci correspond to liquid-liquid phase separated droplets (14), which in this case, were disrupted by HP.

## CONCLUSION

We have shown for the first time that quantitative, high resolution 2-photon imaging modalities can be applied to study the response of live bacterial, archaeal and eukaryotic microbial cells to high hydrostatic pressure. Significant changes in metabolic state and protein interactions were observed for the mesophilic organisms in response to pressure, while the extremophile archaeon exhibited no pressure-induced change in metabolism. The fact that pathogens such as certain strains of *E. coli* evolve tolerance to and even the ability to grow at high pressure in the context of pressure treatment of food products (30–32), underscores the importance of understanding the mechanism underlying these adaptations. Moreover, to the extent that extremophilic organisms can be genetically manipulated, as is the case with *Photobacterium profundum SS9* (25), it will be possible to engineer fluorescent protein and promoter fusions in these organisms, opening up large avenues of investigation. Given the sheer scale of life in the deep biosphere, and our limited knowledge of the molecular mechanisms at play, high-pressure, high-resolution quantitative imaging will be extremely useful for understanding molecular and cellular adaptations to the environments of the deep biosphere.

## Author Contributions

A. C. Bourges – performed research, A. Lazarev – contributed to system design, N. Declerck – interpreted results and wrote the paper, K.L. Rogers – provided samples and assisted with sample preparation, C.A. Royer – designed research, interpreted results, wrote the paper.

## Acknowledgements

We would like to thank Jean-Yves Bouet for providing the *E. coli* strain DLT3053 with the plasmid pJYB234. This research was funded in part by the Alfred P. Sloan Foundation through the Deep Life Community of the Deep Carbon Observatory.

## SUPPLEMENTAL INFORMATION

**FIGURE S1.**
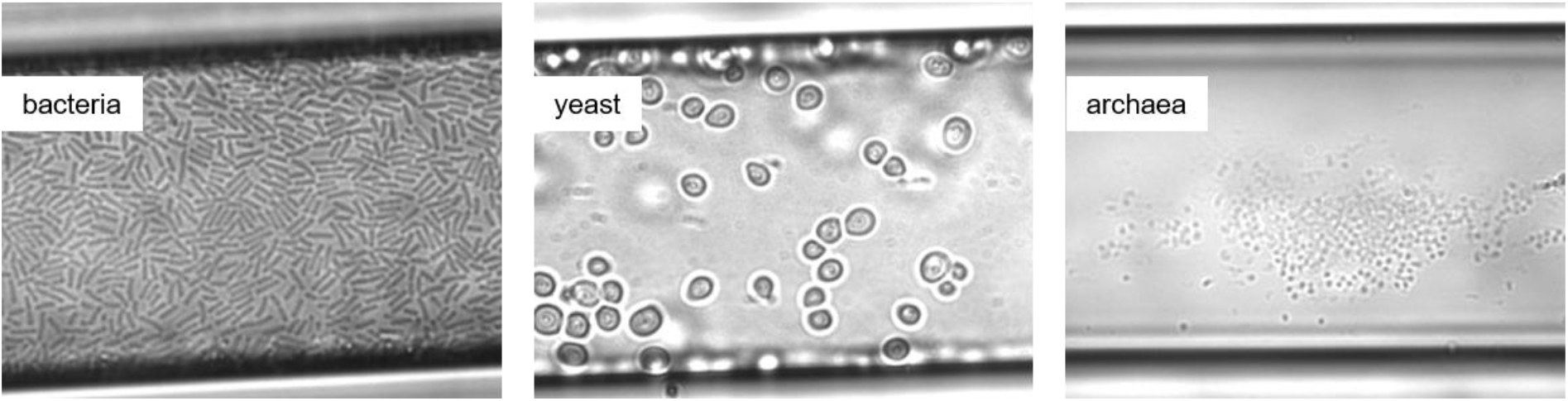
Cells immobilized in the capillary coated with chitosan. Brightfield images of bacteria (*E. coli* MG1655), yeast (*Saccharomyces cerevisiae*) and archaea (*Archaeoglobus fulgidus*) immobilized in a square capillary at atmospheric pressure. The fused silica capillary has an inner diameter of 50 μm and is coated with a chitosan solution prior injection of a highly concentrated cell preparation and washing of the unattached cells.

**FIGURE S2.**
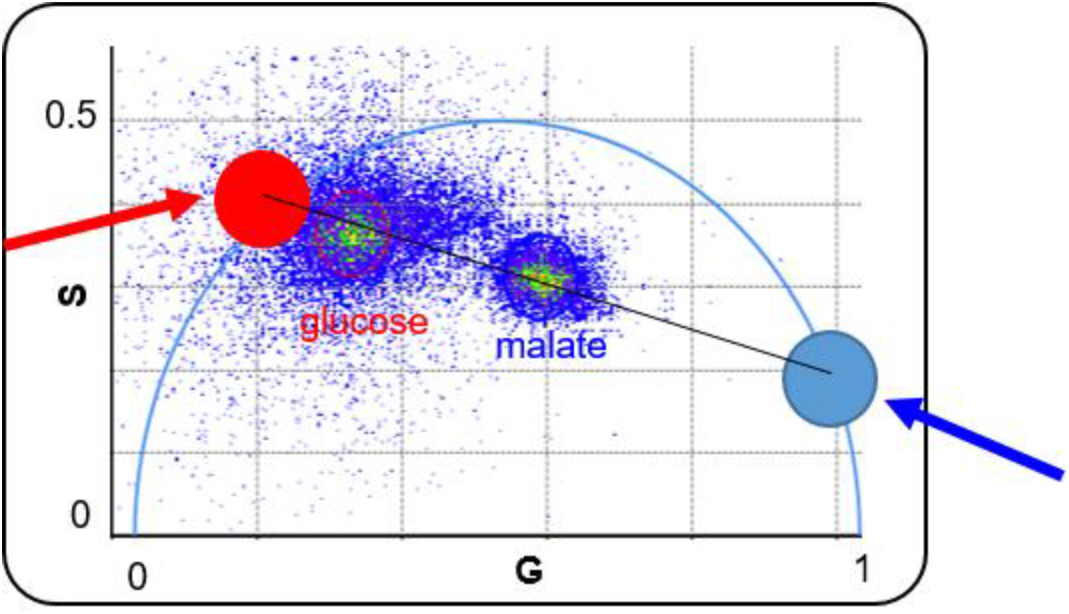
Effect of metabolism of NAD(P)H lifetime. FLIM Phasor analysis of *B. subtilis* NAD(P)H auto-fluorescence grown in media containing a glycolytic carbon source (glucose – red circle) and a gluconeogenic carbon source (malate – blue circle). Arrows indicate the phasor positions on the universal circle at this frequency (80 MHz) of the single exponential decays for free NAD(P)H (blue arrow) and bound NAD(P)H (red arrow). Excitation wavelength was 740 nm.

